# Uniformly processed transcriptome-wide alternative splicing profiles for pediatric cancer research

**DOI:** 10.64898/2026.09.14.750241

**Authors:** Cindy E Liang, Joshua A. Shapiro, Holly C. Beale, Jaclyn N. Taroni, Olena M. Vaske

## Abstract

Alterations in regulatory processes like alternative splicing contribute to pediatric cancer development. Although splicing aberrations have been observed in pediatric leukemias, alternative splicing has yet to be studied in pediatric cancers at scale, due to a lack of uniformly processed, sample-level pediatric cancer splicing profiles with non-diseased tissue comparators. We address this need by quantifying splice event usage for a curated set of bulk RNA-seq datasets from the NCI’s Therapeutically Applicable Research to Generate Effective Treatments (TARGET, n = 1152) and Genotype-Tissue Expression (GTEx, n = 1098) as a comparator. This Treehouse Splice Compendium is accompanied by a reproducible workflow that was used to generate the data in the compendium and reflects the largest known RNA-seq dataset processed by the splice quantification tool Shiba. The compendium is part of a suite of large, uniformly processed datasets aggregated by the UCSC Treehouse Childhood Cancer Initiative and Alex’s Lemonade Stand Foundation’s Childhood Cancer Data Lab, which include the Treehouse Expression Compendia, refine.bio, and the Single-cell Pediatric Cancer Atlas.

## Introduction

High throughput sequencing approaches, most commonly using the Illumina technology, have revolutionized the study of RNA splicing. Bioinformatic analyses of short read RNA sequencing data have catalogued alternative splicing profiles in adult cancer samples, leading to clinically relevant discoveries^1–4^. Multiple splicing quantification tools measure changes in splicing via percent spliced in values (PSI) for transcript features, such as the inclusion of exons and introns in isoforms^1,3,5–7^. PSI values range between 0 and 1 and describe the percent inclusion of a splice event, where an inclusion event represents a transcript isoform that includes a specific splice feature such as an exon or retained intron, and its corresponding exclusion event represents the transcript isoform with that feature spliced out. In splice event-based quantification approaches, a splice event is defined by changes in sets of sequencing reads that map to sequential exon-exon or exon-intron junctions. Splicing events are further categorized into splice event types based on the features being alternatively spliced within a transcript. Although the set of splice events measured from sequencing data differs from tool to tool, common categories of splice events include skipped exon, retained intron, alternative first and last exon, and alternative 3’ or 5’ splice sites.

The splice data produced by such bioinformatic tools have made landscape analyses of perturbed splicing across adult cancers possible, revealing that cancer samples exhibit more novel (unannotated) splicing compared to samples from non-cancerous tissues^3^, different cancer types exhibit cancer-type-specific levels of alternative splicing^2,3,8,9^ and cancer-specific splice events are clinically relevant^10–12^. Furthermore, the ability to categorize splice events into types has shown to be useful in understanding the regulatory mechanisms governing perturbed splicing in cancer. For instance, in adult lung adenocarcinomas, the finding of increased skipped exon events in samples with mutated splicing factor U2AF1 allowed for sequence motif analysis in these features, leading to a model for how mutated U2AF1 can globally dysregulate splicing through altered U2AF1 binding preferences ^13^.

Aberrant splicing has also been shown to contribute to pediatric cancer development ^14,15^. Compared to adult cancers, somatic coding sequence mutations in pediatric cancers are less frequent^16^, so it is important for researchers to examine regulatory alterations, such as those found in splicing, to better understand the disease. Pediatric leukemias have been shown to exhibit unique patterns in alternative splicing, compared to their adult counterparts^17^. In addition to using different splice events compared to adult forms of the disease, aberrant splicing in pediatric acute myeloid leukemia has been observed in the absence of mutations in splicing factors^18^. Furthermore, therapeutically targetable splicing alterations have been observed in pediatric brain tumors^19^.

A challenge to studying splicing from pediatric cancer sequence data lies in the relative rarity of pediatric cancer samples compared to adult cancers. Several groups have created large data resources that aim to catalog alternative splicing in cancer and normal tissues. However, there are limitations to applying them to pediatric cancer. Of pan-cancer data resources that produce PSI quantifications of alternative splicing, several lack pediatric datasets^2,4,20^. Other resources include pediatric cancer data and comparator tissue samples, but do not provide sample-level splice quantification values^3^. PSI distributions of alternatively spliced events have been observed to follow bimodal distributions, so a single summary statistic based on the median or mean of a group may mask biological signals that would elucidate disease subtypes^21^. Finally, many existing splice resources do not quantify unannotated splicing events or use tools that perform poorly at unannotated event detection^2^. A summary of the available data resources are provided in Figure 1.

**Fig 1:**
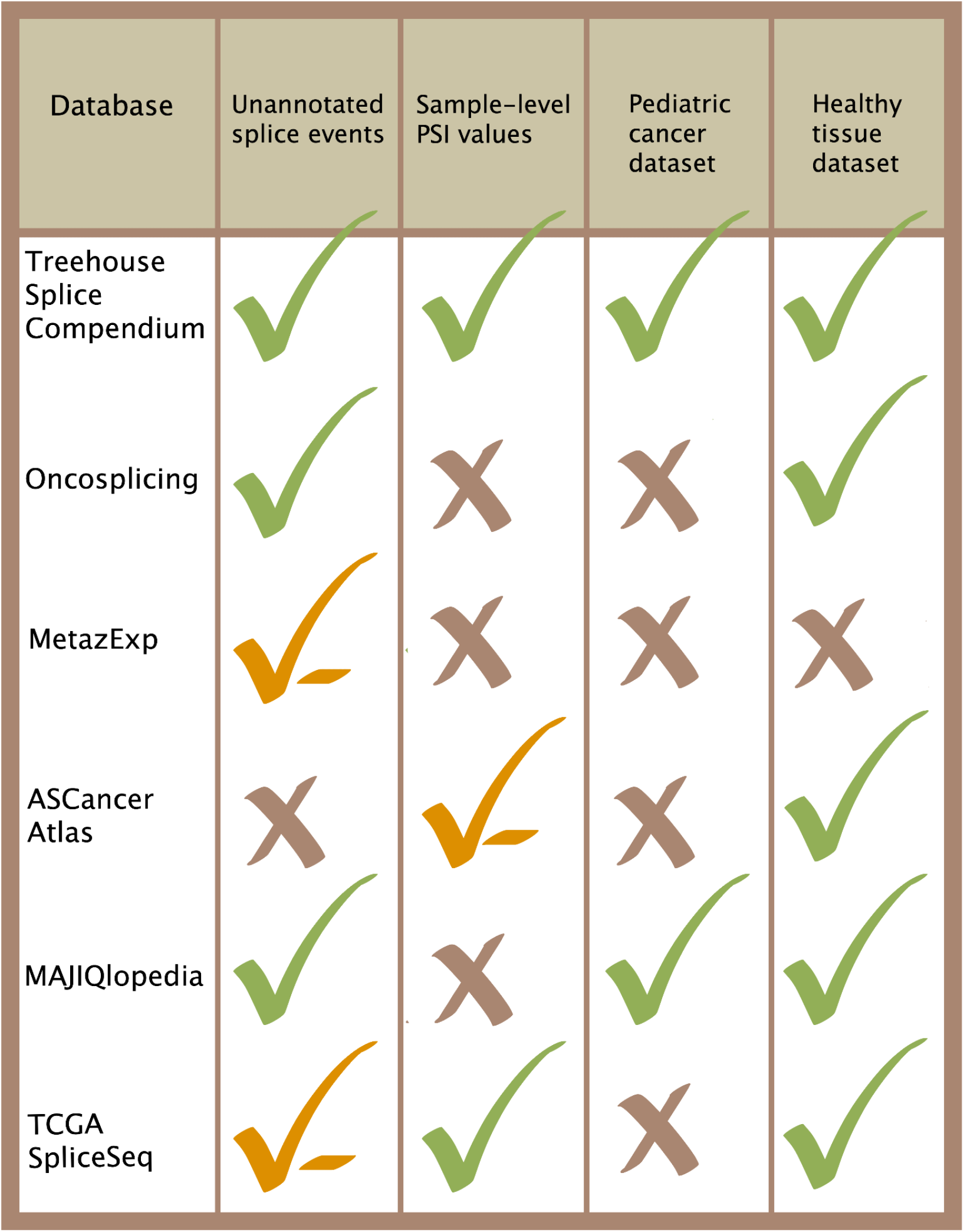
Comparison of existing data resources that contain PSI quantifications of alternative splicing events. Checkmarks indicate the presence of the features in each column. A checkmark-minus is given for the following caveats: MetazExp^22^ quantifies alternative splicing using rMATS, which has performed poorly in unannotated splice event detection in evaluations^6,23^; ASCancerAtlas^20^ does not contain uniformly processed PSI values with the same tool; TCGA SpliceSeq does not quantify events with unannotated exons^2^. Although MAJIQ (used by MAJIQlopedia) is not a true event-based tool, it categorizes event types into “retained intron” or “non retained intron” categories^24^. The “x” indicates that the given feature is absent from the listed data resource.

To address these limitations, we created the Treehouse Splice Compendium, a data resource of uniformly processed pediatric cancer and non-cancer tissue sequence data, to enable researchers to investigate splicing across pediatric cancers and normal tissues. To accomplish this, we used Shiba^6^, an event-based splicing tool that performs well in unannotated event detection.

## Results

We created the Treehouse Splice Compendium (Fig 2) to allow researchers to compare PSI quantifications of annotated and unannotated splice events across pediatric cancer and healthy samples. Given the limitations of existing resources, we prioritized including cancer types and curated healthy tissue comparators for pediatric cancers from large, well-known datasets that are accompanied by clinical information. Version 1 of the Treehouse Splice Compendium consists of a curated set of samples from NCI’s TARGET (Therapeutically Applicable Research to Generate Effective Treatments) initiative and GTEx (Genotype-Tissue Expression) healthy tissue datasets. RNA-Seq datasets from 1152 TARGET samples are included, spanning acute lymphoblastic leukemia (ALL), acute myeloid leukemia (AML), clear cell sarcoma of the kidney (CCSK), rhabdoid tumor (RT), wilms tumor (WT), and neuroblastoma (NBL) cancer types. To serve as healthy tissue comparators for differential analysis, 1098 GTEx samples are included, spanning EBV-transformed lymphocytes, kidney, muscle, and whole blood tissue types (Fig 3). GTEx data are derived from young adult and adult patients, while TARGET data are derived from pediatric patients (Fig 3D). Further demographic and clinical information, such as race/ethnicity and overall survival are provided in the compendium metadata on Zenodo.

**Fig 2:**
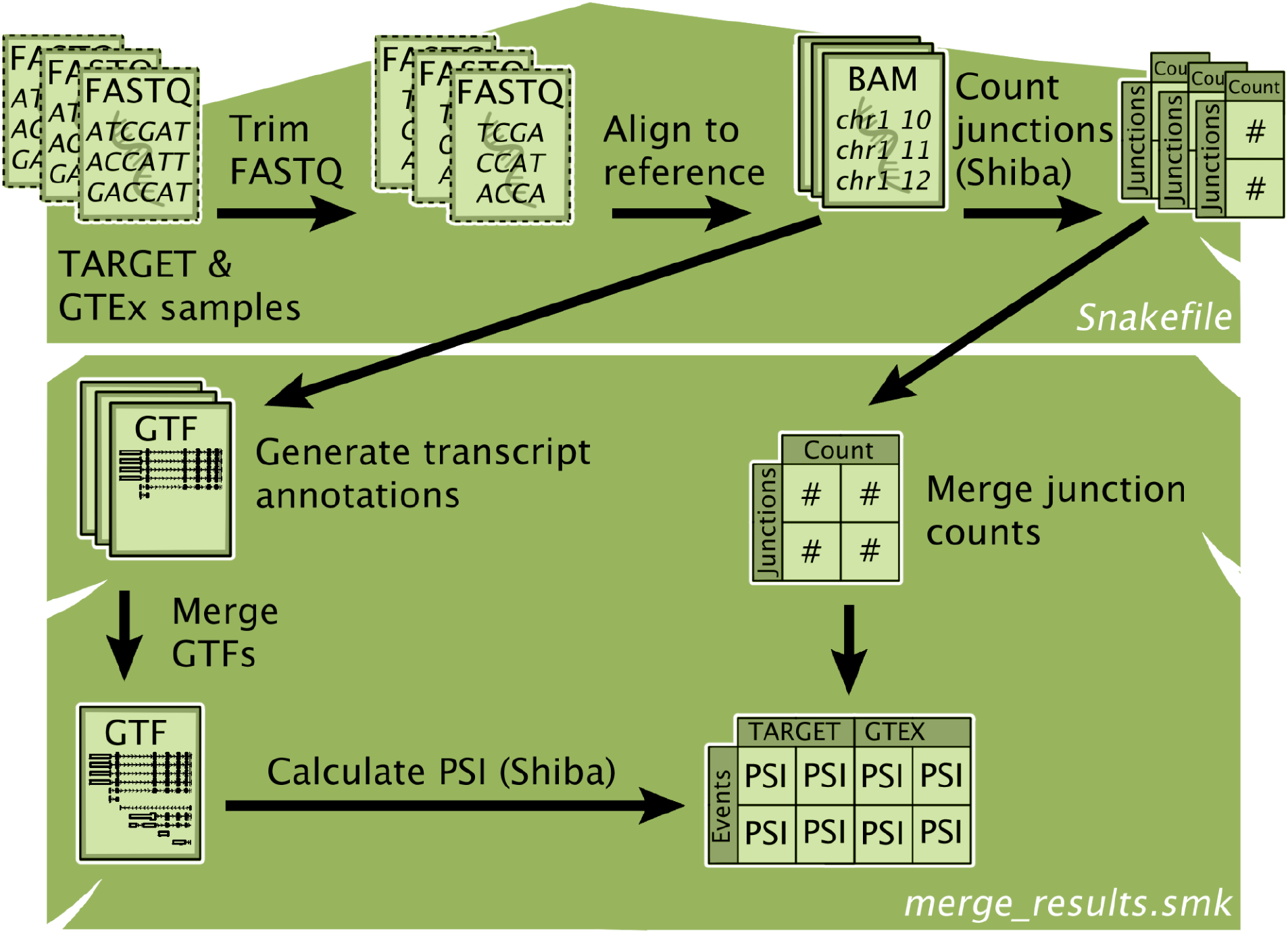
Treehouse Splice Compendium workflow diagram. Data processing is performed using Snakefile and merge_results.smk workflows. Junction count, GTF, and PSI files are produced with Shiba^6^.

**Fig 3:**
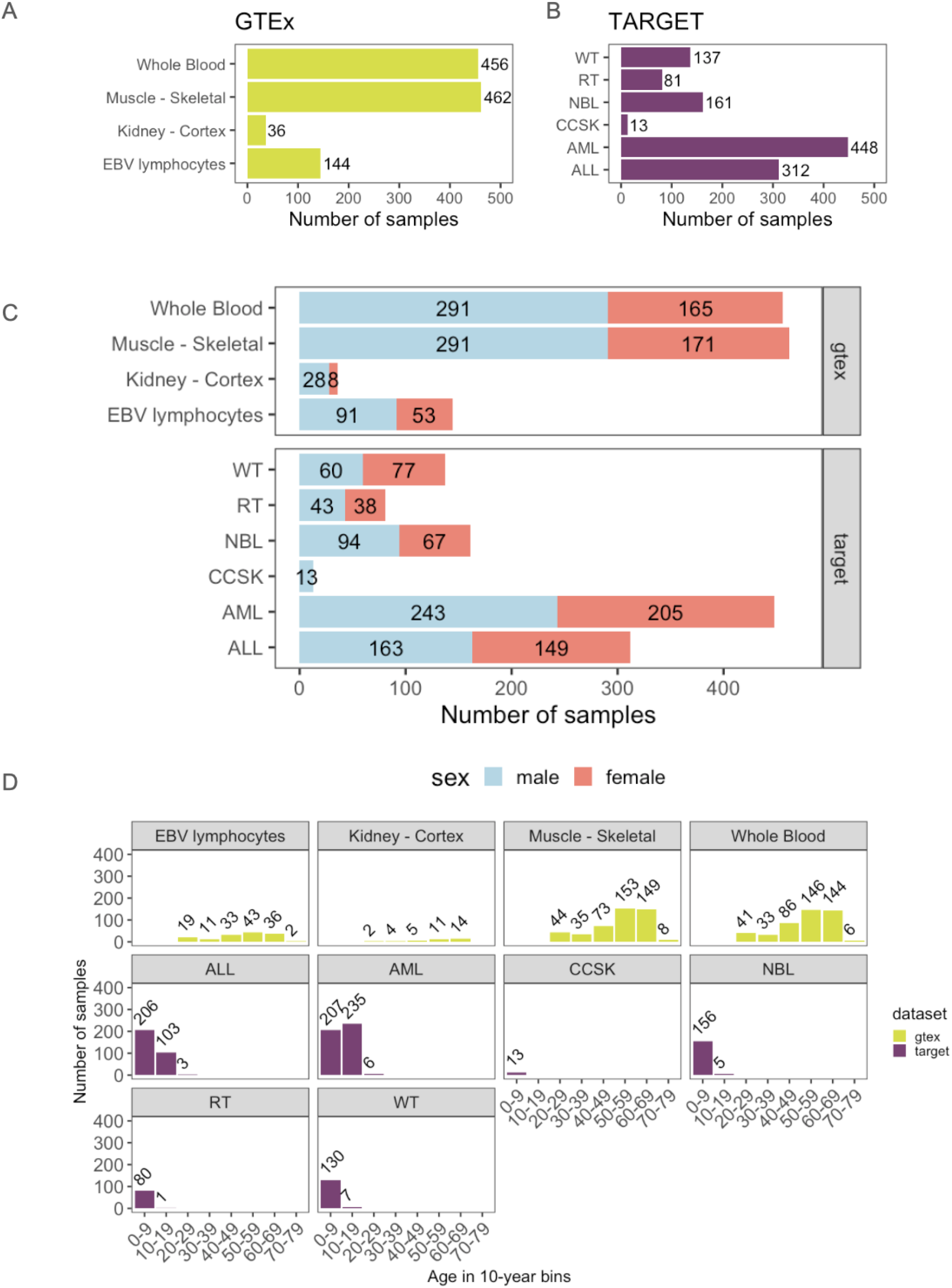
Summary of Treehouse Splice Compendium datasets. A-B: Breakdown of number of samples in each tissue type (y-axis) represented in the Compendium. C: Sex breakdown of Compendium samples, per tissue type. D: Age of donors in 10-year bins, per tissue type. Abbreviations are as follows: EBV lymphocytes refer to Epstein-Barr virus-transformed lymphocyte cell lines from the GTEx dataset. WT - Wilms tumor; RT - Rhabdoid tumor; NBL - Neuroblastoma; CCSK - Clear cell sarcoma of the kidney; AML - Acute myeloid leukemia; ALL - Acute lymphocytic leukemia. TARGET and GTEx data were downloaded from the Sequence Read Archive, with downloads finishing on 5/10/26.

## Methods

### Data description

Splice compendium data consists of the following components: 1) a processed PSI matrix composed of sample-level PSI values with additional fields such as gene name and splice event annotation status for quick code-free filtering and 2) a metadata table providing tissue type, clinical, and demographic information of each sample in the compendium. Additionally, all scripts used to generate these data are provided in a public GitHub repository (see Data Availability).

### Data processing

RNA sequencing datasets were obtained from the [GTEx Portal](https://gtexportal.org/home/) on 04/29/2026 and bGaP accession number phs000424 on 05/10/2026 and the Therapeutically Applicable Research to Generate Effective Treatments (https://www.cancer.gov/ccg/research/genome-sequencing/target) initiative, phs000218. A subsample of these datasets were used, based on the following criteria: total RNA sequence samples that are poly-A selected, paired-end, and falling under the tissue types listed in Figure 3. The full experimental procedure for RNA-seq data generation for these samples are described in the TARGET Cancers Selected for Study page^25^ and the GTEx portal methods page^26^.

Data processing to generate event-based splice quantifications from FASTQ sequence files was performed using the Treehouse Splice Compendium workflow. Briefly, RNA sequence files were downloaded as FASTQs using SRA-toolkit tools^27^. FASTQs were trimmed with FastP^28^ prior to alignment with STAR^28,29^. Aligned files were sorted and indexed using samtools sort and samtools index^30^ with default parameters.

Alternative splicing is quantified using Percent Spliced-In (PSI) values with Shiba v0.8.1. The default mode of Shiba combines gene annotations from all samples using stringtie --merge to build a comprehensive database of all possible splice events for quantification. To take advantage of this feature, we generate GTFs from the aligned BAMs of each individual sample with StringTie^31^, using the reference GTF as a guide. Then, GTFs of all samples in the compendium are merged, so that a comprehensive set of transcript annotations can be used for defining splice events. The merged GTF is passed into Shiba’s events2gtf.py to generate coordinates of splice events.

To generate junction counts for each sample that are used to calculate PSI, we ran Shiba’s bam2junc.py on each BAM file in the compendium. These per-sample junctions were then merged to create a single junction file, which was used as input alongside the event coordinates to calculate PSI with Shiba’s psi.py.

To our knowledge, the Treehouse Splice Compendium represents the largest released dataset of sample-level PSI values that has been processed with Shiba to date. In order to process a dataset this size, we split processing into a number of functions described in the workflow. Results of our split workflow are identical to the canonical Shiba approach for all events, except retained introns, which we therefore do not include in this release. For samples in this compendium, PSI values are calculated for a splice event if the event’s junctions have at least 10 reads. Otherwise, no PSI value is given for that sample in the matrix.

## Discussion

The Treehouse Splice Compendium is a data resource that meets the need for uniformly processed pediatric cancer-focused splice data by providing sample-level PSI values of annotated and unannotated alternative splice events. RNA-seq analyzed in this compendium span six pediatric cancer types and four undiseased tissue types. By providing sample-specific splice quantifications, we anticipate that this resource will facilitate studies related to understanding alternative splicing-driven heterogeneity and subtypes within pediatric cancers. Researchers who study specific alternative splicing events in other contexts, such as in adult cancers, may also be interested in understanding the range of diseases that exhibit splicing for a given splice event, defined by any combination of event type, gene, and chromosome coordinate information recorded in the released PSI matrix. We anticipate that this resource will facilitate hypothesis-generating research from experts in the adult cancer splicing space, who have yet to explore splicing in pediatric cancers.

Future releases of the Treehouse Splice Compendium are planned to expand the number and type of data, as well as to address two limitations. We will incorporate retained intron event quantifications using other splice quantification tools such as MAJIQ and incorporate samples from the developmental GTEx project so there can be more representation from age-matched healthy reference groups.

## Acknowledgements

The results published here are in whole or part based upon data generated by the Therapeutically Applicable Research to Generate Effective Treatments (https://www.cancer.gov/ccg/research/genome-sequencing/target) initiative, phs000218. (https://portal.gdc.cancer.gov) and the Genotype-Tissue Expression (GTEx) Project.

## Supplemental

**Supp table 1.** Number of GTEx samples, categorized by tissue type and sequencing center of origin.

| tissue_type | center_name | n |
| --- | --- | --- |
| Muscle - Skeletal | Broad Institute | 462 |
| Whole Blood | Broad Institute | 456 |
| Cells - EBV-transformed lymphocytes | Broad Institute | 144 |
| Kidney - Cortex | Broad Institute | 36 |

**Supp Table 2.** Number of TARGET samples, categorized by cancer type and sequencing center of origin. BCCAGSC = British Columbia Cancer Agency Genome Sciences Centre; STJUDE = St. Jude Children’s Research Hospital; HAIB = HudsonAlpha Institute for Biotechnology; NCI-KHAN = National Cancer Institute.

| study_name | Center Name | n |
| --- | --- | --- |
| TARGET: Acute Lymphoblastic Leukemia (ALL) Expansion Phase 2 | BCCAGSC | 304 |
| TARGET: Acute Lymphoblastic Leukemia (ALL) Pilot Phase 1 | STJUDE | 3 |
| TARGET: Acute Myeloid Leukemia (AML) | BCCAGSC | 371 |
| TARGET: Acute Myeloid Leukemia (AML) | HAIB | 64 |
| TARGET: Kidney, Rhabdoid Tumor (RT) | BCCAGSC | 66 |
| TARGET: Kidney, Wilms Tumor (WT) | BCCAGSC | 122 |
| TARGET: Neuroblastoma (NBL) | NCI-KHAN | 146 |

**Supp Table 3.** Number of GTEx samples, categorized by tissue type and the version release they originate from.

| tissue_type | version | n |
| --- | --- | --- |
| Muscle - Skeletal | 2 | 449 |
| Whole Blood | 2 | 429 |
| Cells - EBV-transformed lymphocytes | 2 | 131 |
| Kidney - Cortex | 3 | 2 |
| Kidney - Cortex | 2 | 34 |
| Whole Blood | 3 | 17 |
| Cells - EBV-transformed lymphocytes | 3 | 5 |
| Muscle - Skeletal | 3 | 11 |
| Whole Blood | 1 | 10 |
| Cells - EBV-transformed lymphocytes | 1 | 8 |
| Muscle - Skeletal | 1 | 2 |

**Supp table 4.** Number of TARGET samples, categorized by cancer type and the version release they originate from.

| tissue_type | version | n |
| --- | --- | --- |
| Acute Lymphoblastic Leukemia (ALL) Expansion Phase 2 | 1 | 307 |
| Acute Lymphoblastic Leukemia (ALL) Expansion Phase 2 | 2 | 2 |
| Acute Lymphoblastic Leukemia (ALL) Pilot Phase 1 | 1 | 3 |
| Acute Myeloid Leukemia (AML) | 1 | 448 |
| Kidney, Clear Cell Sarcoma of the Kidney (CCSK) | 1 | 13 |
| Kidney, Wilms Tumor (WT) | 1 | 137 |
| Kidney, Rhabdoid Tumor (RT) | 1 | 81 |
| Neuroblastoma (NBL) | 3 | 32 |
| Neuroblastoma (NBL) | 1 | 128 |
| Neuroblastoma (NBL) | 2 | 1 |

## Code availability

Code used to process data in the Treehouse Splice Compendium are found in the following github repo: https://github.com/UCSC-Treehouse/splicing-compendium

## Data availability

The data used for the analyses described in this manuscript were obtained from: the [GTEx Portal](https://gtexportal.org/home/) on 04/29/2026 and [dbGaP](http://www.ncbi.nlm.nih.gov/gap) accession number phs000424 on 05/10/2026; the Therapeutically Applicable Research to Generate Effective Treatments (https://www.cancer.gov/ccg/research/genome-sequencing/target) initiative, phs000218. The TARGET clinical metadata used for this analysis are available at the Genomic Data Commons (https://portal.gdc.cancer.gov) and UCSC Xena^32^.

A data table of PSI values quantified for samples in the compendium, alongside a clinical metadata table with survival and patient demographic information, can be downloaded at https://zenodo.org/records/22737043.

## Declarations

J.A.S. and J.N.T. are employees of Alex’s Lemonade Stand Foundation, a funder of this work.

